# *miR-263b* controls circadian locomotor activity and the structural plasticity of small ventral lateral neurons by inhibition of Beadex

**DOI:** 10.1101/368043

**Authors:** Xiaoge Nian, Wenfeng Chen, Weiwei Bai, Zhangwu Zhao, Yong Zhang

## Abstract

Circadian clocks drive rhythmic physiology and behavior to allow adaption to daily environmental changes. In *Drosophila*, the small ventral lateral neurons (sLNvs) are the master pacemakers that control circadian rhythms. Circadian changes are observed in the dorsal axonal projections of the sLNvs, but their physiological importance and the underlying mechanism are unclear. Here we identified *miR-263b* as an important regulator of circadian rhythms in *Drosophila*. Flies depleted of *miR-263b* (*miR-263b^KO^*) exhibited dramatically impaired rhythms under constant darkness. Indeed, *miR-263b* is rhythmically expressed and controls circadian output by affecting the structural plasticity of sLNvs through inhibition of expression of the LIM-only protein *Beadex (Bx)*. The misexpression of *Bx* in flies phenocopied *miR-263b^KO^* in behavior and molecular characteristics. In addition, the circadian phenotypes of *miR-263b^KO^* were recapitulated by mutating the *miR-263b* binding sites in the *Bx* 3’UTR. Together, these results establish *miR-263b* as an important regulator of circadian locomotor behavior.

## Introduction

Circadian clocks are intracellular pacemakers that generate approximately 24-hour rhythms of behavior and physiology in most organisms. In animals, the core circadian oscillator consists of a conserved autoregulatory transcriptional-translational negative feedback loop (Crane & Young, 2014; Hardin & Panda, 2013; Tataroglu & Emery, 2015). In *Drosophila*, transcription factors CLOCK (CLK) and CYCLE (CYC) activate rhythmic transcription of clock-controlled genes. Among these genes, PERIOD (PER) is a key repressor in the core circadian negative feedback loop that inhibits CLK/CYC activity and represses *per* transcription. Post-translational modifications such as phosphorylation, glycosylation, and ubiquitination also play important roles in setting the pace of circadian clock (J. C. Chiu, Ko, & Edery, 2011; Grima, Dognon, Lamouroux, Chelot, & Rouyer, 2012; Hardin & Panda, 2013; Kim et al., 2012; Ko, Jiang, & Edery, 2002; W. F. Luo et al., 2012).

Circadian locomotor rhythms of flies are generated by a neuronal network consisting of about 150 circadian neurons in the brain (Beckwith & Ceriani, 2015; Dubowy & Sehgal, 2017; Johard et al., 2009; Nitabach & Taghert, 2008). Based on location and function, these circadian neurons are divided into clusters: three groups of ventral lateral neurons (LNvs), three groups of dorsal neurons (DN1s, DN2s, and DN3s), dorsal lateral neurons (LNds), and lateral posterior neurons (LPNs). The large LNvs (lLNvs) and four of the small LNvs (sLNvs) express the neuropeptide PDF, whereas the fifth sLNv is PDF negative (Nitabach & Taghert, 2008). Flies exhibit bimodal locomotor activity rhythms, peaking around dawn and dusk. PDF-positive sLNvs are responsible for promoting the morning peak before daylight, and the fifth sLNv and the LNds are mainly responsible for generating the evening activity (Grima, Chelot, Xia, & Rouyer, 2004; Stoleru, Peng, Agosto, & Rosbash, 2004). DN1s can integrate light or temperature inputs and are able to generate either morning or evening activity; DN1s also promote sleep (Guo et al., 2016; Zhang, Liu, Bilodeau-Wentworth, Hardin, & Emery, 2010). A recent study indicated that the DN1 activity is acutely modulated by temperature, thus DN1s sense temperature to regulate sleep (Yadlapalli et al., 2018). On the other hand, DN2s play important roles in the temperature preference of circadian rhythm (Kaneko et al., 2012).

As master pacemaker neurons, the PDF-positive sLNvs synchronize other circadian neurons, thus enabling flies to maintain robust circadian locomotor rhythms in constant conditions (Renn, Park, Rosbash, Hall, & Taghert, 1999). Elimination of sLNvs or mutation of *pdf* or its receptor causes arrhythmic behavior (Mertens et al., 2005; Renn et al., 1999). The PDF positive sLNvs also send axonal projections toward the dorsal protocerebrum region of fly brain, where DN1s and DN2s are located (Helfrichforster, 1995; Helfrichforster & Homberg, 1993). PDF immunoreactivity in the dorsal axonal terminal of sLNvs has a clearly circadian pattern with high intensity during the daytime and low intensity at nighttime (Park et al., 2000). The sLNvs dorsal projections also show arborization rhythms with higher complexity of axon terminals found in the early day and lower complexity at the night (Fernandez, Berni, & Ceriani, 2008). This structural plasticity of the sLNvs dorsal projections is controlled by both circadian and activity dependent mechanisms (Depetris-Chauvin et al., 2011; Muraro, Pirez, & Ceriani, 2013). The molecular mechanisms underlying this plasticity of sLNv are poorly understood, however. The transcription factor Mef-2 was found to control circadian plasticity of sLNvs through regulation of the cell adhesion molecule Fas2 (Sivachenko, Li, Abruzzi, & Rosbash, 2013). In addition, two matrix metalloproteinases (MMP1, and MMP2) are also shown to be required for the structural remodeling control of sLNv projections (Depetris-Chauvin et al., 2014).

Recently the circadian protein Vrille was found to control sLNv arborization rhythms through unknown mechanisms (Gunawardhana & Hardin, 2017). microRNAs (miRNAs) are small non-coding RNAs of about 22 nucleotides that regulate gene expression post-transcriptionally (Bartel, 2004). miRNAs repress expression of their target genes through mRNA degradation and/or translation inhibition. Recent studies have uncovered critical roles of miRNAs in the regulation of different aspects of animal circadian clocks (Mendoza-Viveros et al., 2017; Xue & Zhang, 2018). In mouse, *miR-132/212* modulates circadian entrainment to day length, thus influencing seasonal adaptation, while *miR-219* and *miR-24* affect circadian period length (Cheng et al., 2007; Yoo et al., 2017). In *Drosophila*, overexpression of the miRNA *bantam* lengthens circadian locomotor period by repressing *clk*, and *timeless* and *clockwork orange* are targets of *miR-276a* and *let-7*, respectively (W. F. Chen et al., 2014; X. Chen & Rosbash, 2016; Kadener et al., 2009). miRNAs also play important roles in circadian output pathways. We showed previously that the disruption of general miRNA functions resulting from depletion of GW182 affects circadian rhythms by interfering with PDF-receptor signaling (Zhang & Emery, 2013). In addition, the *miR-279* and *miR959–964* cluster miRNAs modulate circadian locomotor behavior output and circadian timing of feeding and immune responses (W. Y. Luo & Sehgal, 2012; Vodala et al., 2012). Others and we have demonstrated that *miR-124* controls the phase of circadian locomotor activity (Garaulet et al., 2016; Zhang, Lamba, Guo, & Emery, 2016).

To further understand the post-transcriptional regulation of circadian clock functions by miRNAs, we screened for circadian clock phenotypes in miRNA mutants. Here we show that a conserved miRNA, *miR-263b*, is an important regulator of circadian rhythms in *Drosophila*. *miR-263b* regulates circadian locomotor behavior by affecting the structural plasticity of sLNvs through inhibition of Beadex (Bx) expression.

## Results

### *miR-263b* regulates circadian behavior

To identify miRNAs that regulate circadian rhythms, we first used miRNA quantitative real-time PCR to identify rhythmically expressed miRNAs in fly heads. We found that the expression of *miR-263b* is rhythmic in fly heads, consistent with previous microarray data, (Figure 1A, (Yang, Lee, Padgett, & Edery, 2008)). Importantly, the oscillation of *miR-263b* was abolished in the *Clk^Jrk^* mutant confirming that the expression of *miR-263b* is under circadian control (Figure 1A).

**Figure 1.**
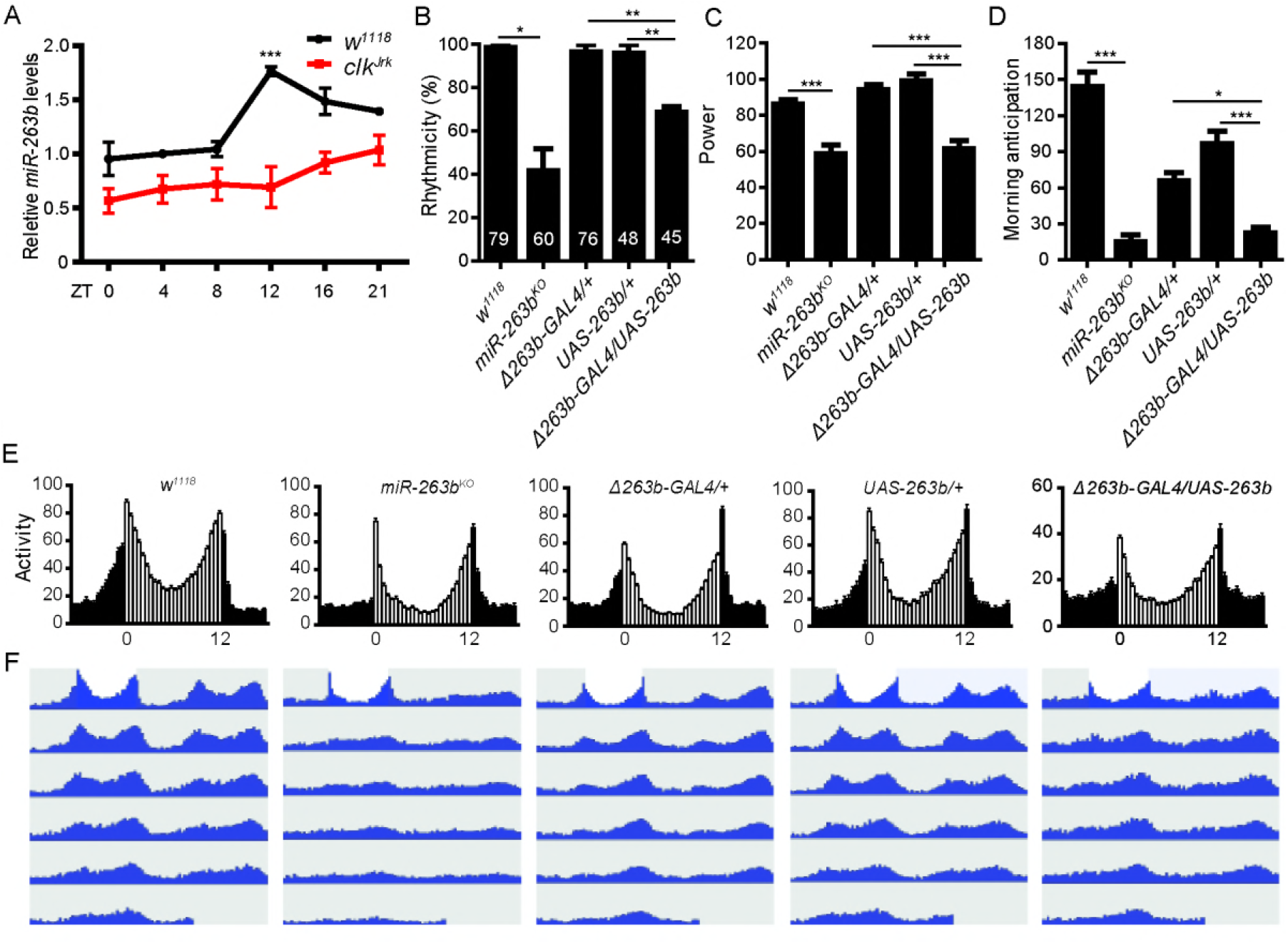
*miR-263b* is rhythmically expressed and regulates circadian behavior. Quantitative-PCR analysis of *miR-263b* from adult brains of wild-type and *Clk^Jrk^* mutant flies at the indicated time points. The relative expression levels were normalized to 2S rRNA and were further normalized to *w^1118^* at ZT0. Each time point was compared to ZT0. Data represent mean ± SEM. ***P< 0.001 determined by Student’s t test. (B-D) Comparison of B) rhythmicity, C) power, and D) morning anticipation index in indicated fly strains. Data represent mean ± SEM (n=45–79). *P<0.05, ^**^P<0.01, ***P< 0.001 determined by Student’s t test. (E) Locomotor activity of indicated strains measured during 3 days of LD. The white and black bars indicate day and night, respectively. (F) Actograms showing the average activities on the last day in LD and 5 days in DD. Light represents the day and gray darkness. See also Figure S1.

To examine the circadian role of *miR-263b*, we analyzed the previously generated *miR-263b* knockout (*miR-263b^KO^*) flies (Hilgers, Bushati, & Cohen, 2010). In constant darkness (DD) condition, flies lacking *miR-263b* displayed a severely disrupted circadian locomotor rhythm (Figure 1B, Table 1). In contrast to the wild-type flies, more than 60% of the *miR-263b^KO^* flies were arrhythmic (Figure 1B). Moreover, the amplitude of behavioral rhythms was clearly reduced even in rhythmic *miR-263b^KO^* flies (Figure 1C). Because of the high percentage of arrhythmia of the *miR-263b^KO^* flies in DD, we examined the locomotor activity under light dark cycles (LD). Under LD, wild-type flies gradually increase their activity before light-on, an anticipatory behavior controlled by the clock. In *miR-263b^KO^* flies, however, the morning anticipation was abolished (Figure 1D and 1E).

**Table 1:** Locomotor activity of flies with altered *miR-263b* and *Bx* levels in DD

|  |  |  |  |  |
| --- | --- | --- | --- | --- |
| Genotype | N | % Rhythmic | Period (hr) ± S.E.M. | Power* ± S.E.M |
| *w*<sup>1118</sup> | 79 | 98.5±1.5 | 24.4±0.1 | 86.0±2.7 |
| *miR-263b*<sup>KO</sup> | 60 | 41.5±10.4 | 24.3±0.1 | 58.7±4.8 |
| *Bx* mutant | 105 | 40.1±4.8 | 25.7±0.2 | 44.7±2.8 |
| Δ263b-GAL4/UAS-263b | 45 | 68.8±2.6 | 24.2±0.2 | 61.4±4.5 |
| UAS-*Bx*/+; Δ263b-GAL4/+ | 48 | 25.1±10.9 | 24.4±0.4 | 48.6±6.3 |
| Δ263b-GAL4/+ | 76 | 96.6±3.3 | 24.5±0.0 | 94.1±3.7 |
| UAS-263b/+ | 48 | 95.8±4.2 | 23.8±0.1 | 99.2±3.7 |
| UAS-*Bx*/+ | 45 | 100.0±0.0 | 24.0±0.1 | 105.7±3.6 |
| *Bx* *1 | 71 | 43.6±11.7 | 24.1±0.1 | 69.5±4.3 |
| *Bx* *2 | 69 | 31.1±6.7 | 24.3±0.1 | 54.3±5.4 |
| *yw*, *Act5C-Cas9* | 48 | 100.0±0.0 | 24.4±0.2 | 89.1±4.0 |

\*Power is a measure of rhythm amplitude and corresponds to the height of the periodogram peak above the significance line.

Next, we utilized a GAL4 knock-in replacement (*Δ263b-GAL4*, (Hilgers et al., 2010)) to drive expression of *miR-263b*, resulting in a 9-fold increase of *miR-263b* abundance (Figure S1). The overexpression of *miR-263b* disrupted circadian locomotor activity rhythms in DD (Figure 1B and 1C). In addition, a significant reduction of morning anticipation was also observed in these flies (Figure 1D and 1E). Together, these results indicate that *miR-263b^KO^* and overexpression of *miR-263b* both decrease the robustness of circadian locomotor activity rhythms and suppress the morning anticipation in LD. Notably, the free-running period was not significantly altered in the flies in which *miR-263b* levels were abnormal (Figure 1F, Table1).

### *miR-263b* is required for the circadian structural plasticity of sLNv axonal projections

To examine whether the impaired circadian locomotor rhythmicity in *miR-263b^KO^* flies is due to a defect in the molecular pacemaker or in the circadian output pathway, we examined the oscillation of the key pacemaker protein PER in DD in three important groups of circadian neurons: sLNvs, LNds, and DN1s. We found no obvious changes in PER oscillation in these neurons known to control locomotor behavior (Figure S2), suggesting that *miR-263b* may regulate circadian behavior by modulating the output pathway.

PDF-positive sLNvs send axonal projections toward the dorsal protocerebrum region (Helfrichforster & Homberg, 1993). In addition, circadian structural remodeling of sLNv dorsal projections has been observed (Fernandez et al., 2008). To determine whether *miR-263b* affects the circadian plasticity of PDF-positive projections, we examined the termini of sLNv dorsal projections in *miR-263^KO^* flies with a PDF-specific antibody at early day (Zeitgeber time 2 (ZT2), ZT0 is light on and ZT12 is light off) and early night (ZT14). Similar to previous reports, we found that the wild-type flies had more branches of axon terminals at ZT2 than at ZT14 (Figure 2A and 2B). The axonal crosses were quantified with Sholl’s analysis to assay the interactions between axon branches and concentric circles (see Methods, (Sivachenko et al., 2013)). Remarkably, this difference was abolished in the *miR-263b^KO^* flies due to a significant decrease of axonal crosses at ZT2 compared to control *w*^1118^ flies (Figure 2B and 2C). No significant changes of PDF levels were detected in the sLNv soma, indicating that the decrease of axonal branches was not due to the decrease of PDF staining (Figure 2A, Figure S3).

**Figure 2.**
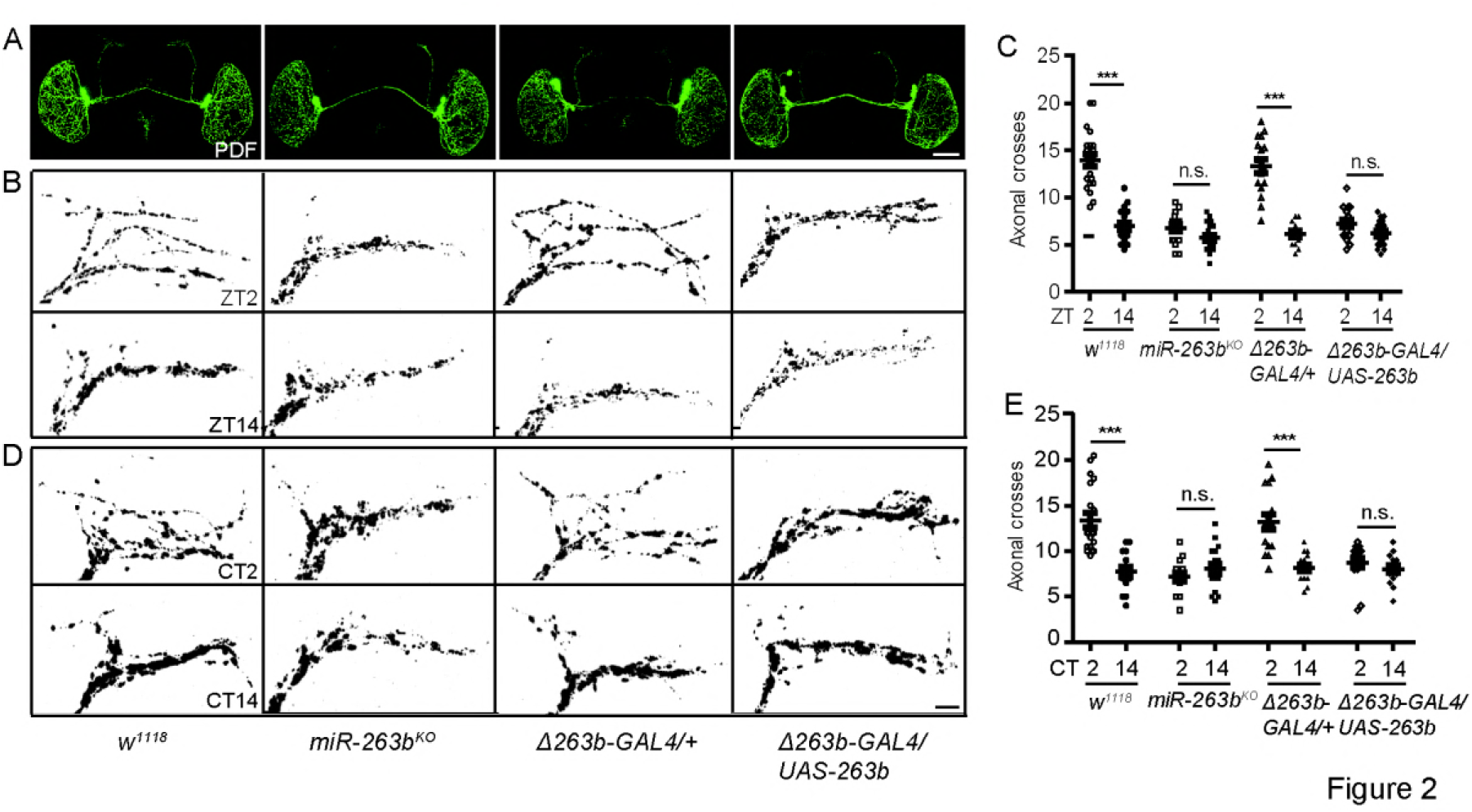
*miR-263b* modulates the structural plasticity of sLNv axonal projections. (A) Representative confocal images of whole brains from the indicated genotypes stained for PDF at ZT2. Scale bar, 75 µm. (B) Representative images of sLNv dorsal projections from the indicated genotypes stained with anti-PDF at ZT2 and ZT14. Flies were entrained for at least 3 days under LD conditions prior to the assay. Note “open”, defasciculated axonal conformation at ZT2 and fasciculated axons at ZT14 in control *w^1118^* flies. Scale bar, 10 µm. (C) Quantification of axonal morphology (fasciculation) of sLNv dorsal termini in LD conditions by Sholl’s analysis (see also STAR* Methods). Data represent means ± SEM (n =15–22). n.s. no significant, ***P< 0.001 determined by Student’s t test. (D) Representative images of sLNv dorsal projections from the indicated genotypes stained with anti-PDF at CT2 or CT14 during the second day of DD. Scale bar, 10 µm. (E) Quantification analysis of axonal morphology of sLNv dorsal termini in DD conditions. Data represent means ± SEM (n =15–21). n.s. no significant, ***P< 0.001 determined by Student’s t test.

To test whether sLNv dorsal projections were present in the absence of *miR-263b*, a membrane-tethered GFP (mCD8-GFP) was used to mark the PDF projections under the control of a *pdf*-specific promoter. As observed by PDF staining, this marker revealed that the dorsal axonal branches of sLNvs were dramatically reduced (Figure S4). Interestingly, overexpression of *miR-263b* caused similar defects in PDF axonal projections (Figure 2B and 2C). These data indicate that the proper level of *miR-263b* is required for the circadian structural plasticity of sLNv dorsal projections.

The structural plasticity of sLNv dorsal projections is under circadian regulation and is activity dependent (Fernandez et al., 2008; Gorostiza, Depetris-Chauvin, Frenkel, Pirez, & Ceriani, 2014; Petsakou, Sapsis, & Blau, 2015; Sivachenko et al., 2013). To further confirm *miR-263b* affects the circadian structural plasticity of sLNvs, we dissected fly brains in DD (Figure 2D). In wild-type flies, we observed more axonal branches at circadian time 2 (CT2) in the subjective morning and fewer branches in the subjective night (CT14) (Figure 2D and 2E). Similar to the results in LD, the rhythmic change in structural plasticity was abolished in the *miR-263b^KO^* flies and in flies that overexpressed *miR-263b* in DD (Figure 2D and 2E).

sLNv axonal complexity is modified in response to electrical activity, and adult-specific activation of sLNvs results in increased complexity (Depetris-Chauvin et al., 2011). Thus we examined whether this activity-dependent structural plasticity requires *miR-263b*. We took advantage of the temperature-gated TrpA1 channel to increase the neuronal activity of sLNvs (Hamada et al., 2008). Increasing the environment temperature to 30°C for 2 hours beginning at ZT12 specifically activated sLNvs and caused defasciculation of sLNv dorsal termini at ZT14 in flies expressing TrpA1 (Figure 3A and 3B). Activation of sLNvs by temperature treatment also increased the dorsal projection complexity (Figure 3B and 3C). Effects were dependent on TrpA1 expression and were similar in all genotypes, including *miR-263b^KO^*. Together, these results indicate that *miR-263b* is required for the circadian structural plasticity but dispensable for the activity-dependent rhythms in sLNv dorsal projections.

**Figure 3.**
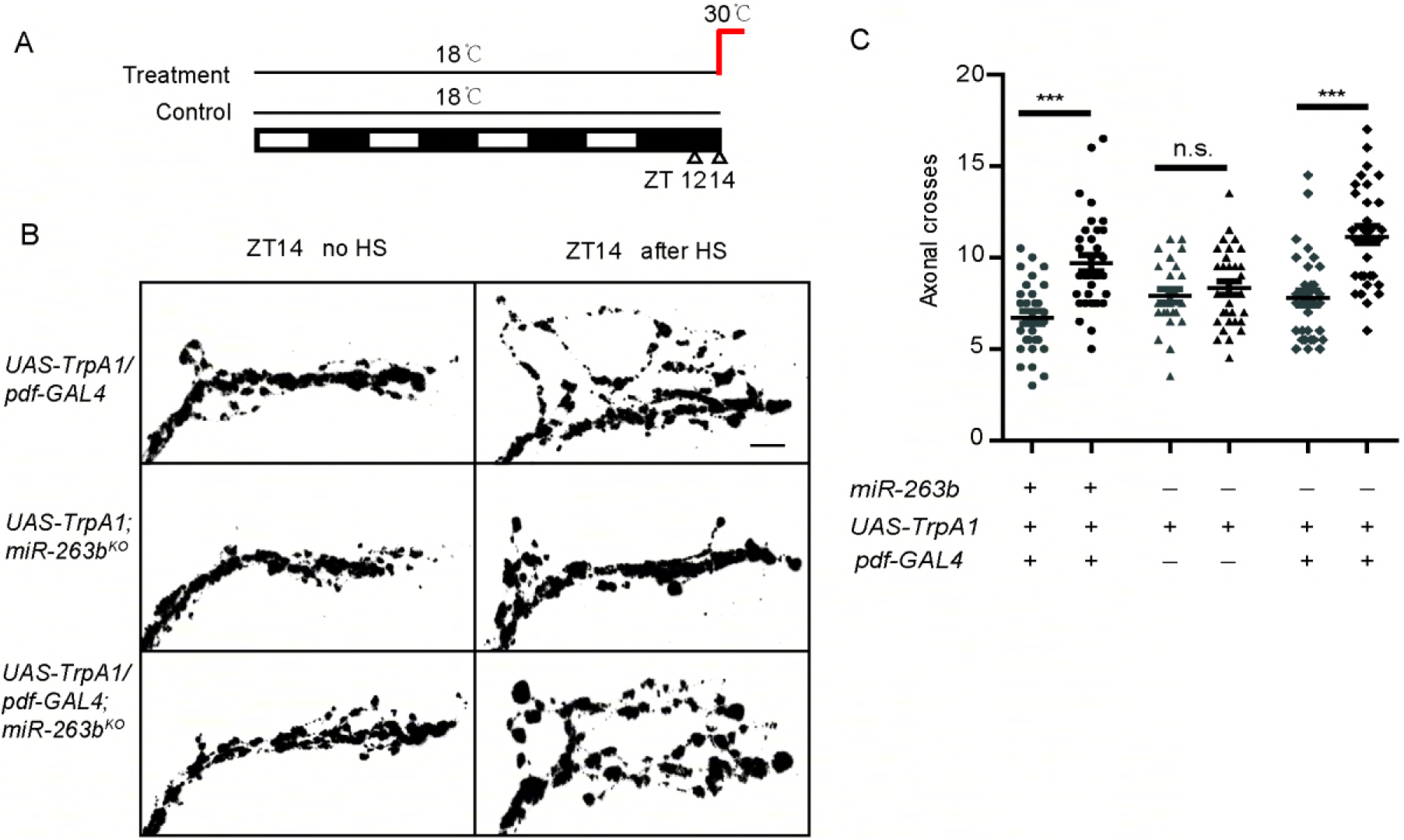
Activity-dependent remodeling of sLNvs axonal fasciculation is intact in *miR-263b^KO^* flies. (A) Diagram of activity induction by heat. Flies were raised at 18°C and entrained in LD cycles at 18°C for at least 3 days before shifting to 30°C at ZT12. Flies were dissected 2hr later (ZT14) for anti-PDF staining. (B) Representative confocal images of sLNvs projections at ZT14 with or without heat shock (HS). Induction of TrpA1 in PDF cells by 2-h temperature elevation to 30°C leads to open conformation of s-LNvs dorsal projections at ZT14 in all strains. Scale bar, 10 μm. (C) Quantification of TrpA1-induced changes in axonal fasciculation. “+” or “-” represents that the relative genes are present or absent in the flies. Data represent means ± SEM (n=26–34). n.s. No significant. ***P< 0.001 determined by Student’s t test.

### Bx regulates circadian rhythms, and its expression is suppressed by *miR-263b*

miRNAs usually regulate gene expression by binding to the 3’ untranslated regions (UTRs) of target mRNAs and causing mRNA degradation and/or translational repression (Bartel, 2004). We used an *in silico* prediction algorithm to identify putative *miR-263b* targets by using the annotated *Drosophila* genome (http://www.targetscan.org/fly_12/).

One of the potential mRNA targets was *Beadex* (*Bx*), which encodes a LIM-only protein. A disrupted circadian rhythm was previously observed in flies with mutations in *Bx* (Tsai, Bainton, Blau, & Heberlein, 2004). Target prediction indicated that the *Bx* 3’UTR has two putative *miR-263b* binding sites, which are highly conserved across *Drosophila* species (Figure S5).

To test whether *miR-263b* directly binds to the *Bx* 3’UTR and inhibits its expression, the *Bx* 3’UTR was fused downstream of a luciferase reporter and transfected into S2 cells. Luciferase activity was significantly suppressed when *miR-263b* was co-transfected with the reporter. In contrast, the luciferase activity was observed in the presence of *miR-263b* when the putative *miR-263b* binding sites within the *Bx* 3’UTR were mutated (Figure 4A). Encouraged by these S2 cell results, we further tested whether Bx abundance is suppressed by *miR-263b* in fly brain. Since the BX antibody we generated in our lab did not work for staining, we decided to use an enhanced GFP (EGFP) reporter strategy. We tested whether *miR-263b* regulates BX levels by expressing an EGFP reporter fused to the wild-type *Bx* 3’UTR or 3’UTR with *miR-263b* binding sites mutated (same construct as Figure 4A). We expressed these two reporters using a *Bx-GAL4* line, which is expressed in the PDF-positive sLNvs and lLNvs, as well as other region of fly brain (Figure S6). Strikingly, we found that the expression of EGFP under the control of the wild-type *Bx* 3’UTR is significantly lower than the mutant 3’UTR in both sLNvs and lLNvs (Figure 4B and 4C). Thus, our S2 cell and imaging results suggest that *Bx* is a direct target of *miR-263b* and is negatively regulated by *miR-263b*. miRNAs are normally negative regulators of gene expression, so we reasoned that overexpression of *Bx* should mimic *miR-263b^KO^*. As expected, overexpression of *Bx* using *Δ263b-GAL4* caused approximately 75% arrhythmicity in DD, which phenocopied the *miR-263b^KO^* (Figure 4D-F, and Table 1). Furthermore, similar to *miR-263b* overexpression, even in the rhythmic *Bx*-mutant flies amplitude of behavior rhythms, and morning anticipation were significantly reduced (Figure 4G and 4H). Taken together, these results demonstrate that Bx is required for robust circadian locomotor rhythms and is a potential target of *miR-263b*.

**Figure 4.**
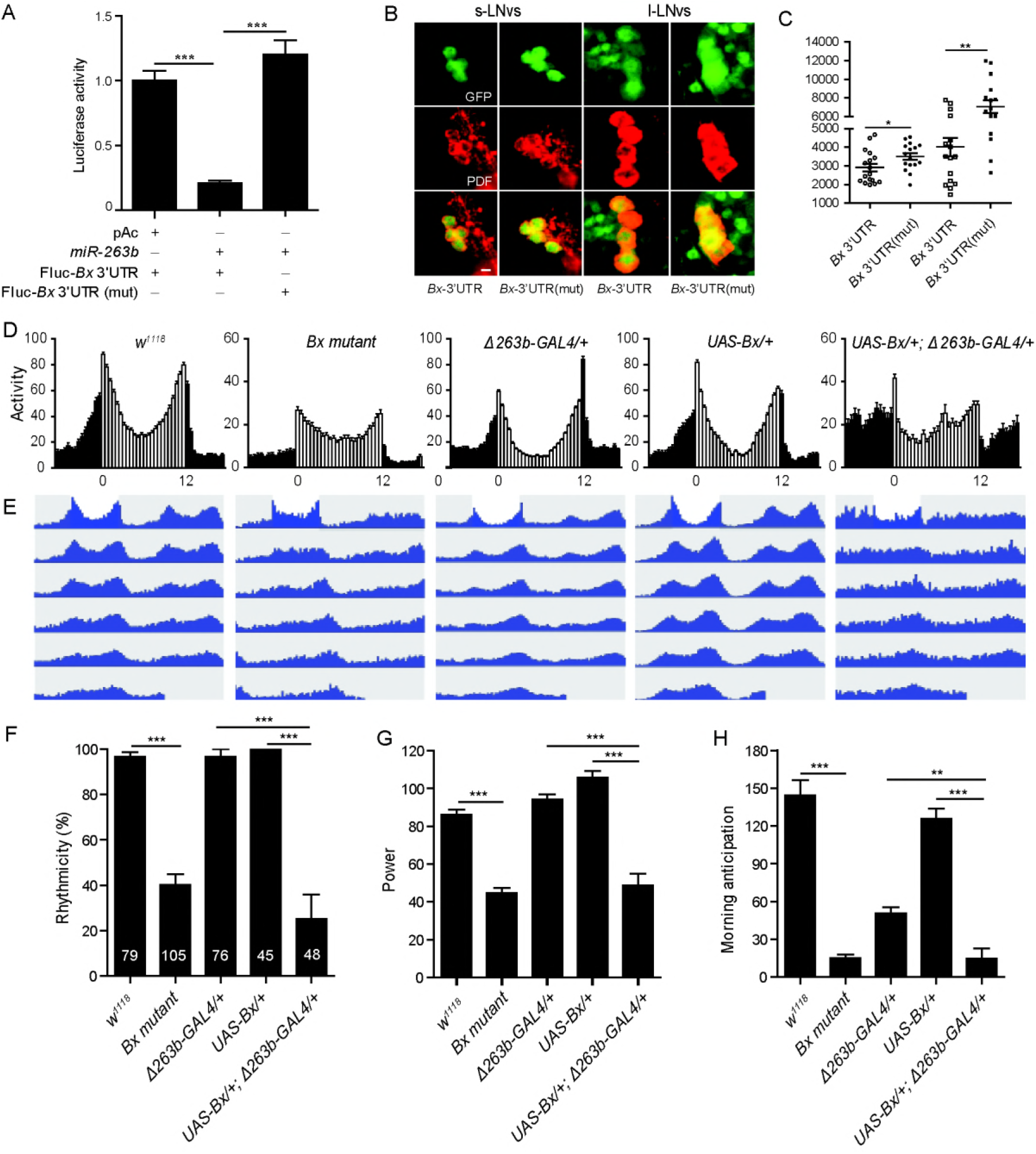
*Bx* regulates circadian rhythms, and its expression is suppressed by *miR-263b*. (A) pAC or pAC-miR-263b were co-transfected with pAc-fluc-Bx 3’UTR or pAc-fluc-Bx 3’UTR (mut) and with pCopia-Renilla luciferase into S2 cells. After two days luciferase activity was quantified. For each condition, a normalized firefly/*Renilla* luciferase value is plotted with SEM. (B) Representative confocal images of *Bx*3’UTR (*Bx-GAL4/+;UAS-EGFP-Bx 3’UTR/+*) and *Bx*3’UTR(mut) (*Bx-GAL4/+;UAS-EGFP-Bx 3’UTR(mut)/+*). The brains were stained with anti-GFP antibody (green) and anti-PDF antibody (red) at ZT13. Scale bar, 10 μm. (C) Quantification of GFP signals in sLNvs and lLNvs from *Bx* 3’UTR and *Bx* 3’UTR(mut) flies. Data represent means ± SEM (n=16–17). *P<0.05, ^**^P<0.01 determined by Student’s t test. (D) Locomotor activity of adult male flies of the indicated genotypes measured during 3 days of LD cycle. The white and black bars indicate day and night, respectively. (E) Actograms showing the average activities on the last day in LD and the 5 days in DD. Light represents the day and gray darkness. (F-H) Comparison of E) rhythmicity, F) power, and G) morning anticipation in *indicated* lines. The numbers of tested flies are shown in each column. Each experiment was conducted three times. Data represent means ± SEM (n=45–105). ^**^P<0.01, ***P< 0.001 determined by Student’s t test.

### Bx regulates the arborization rhythms of sLNv dorsal projections

If *miR-263b* regulates circadian rhythms by downregulating *Bx* expression, the *Bx* mutant should phenocopy the circadian structural plasticity defects in *miR-263b* overexpression. We therefore examined the dorsal axonal projections of sLNvs in *Bx* mutant flies. As expected, PDF projections were maintained in the fasciculated state during the day as well as the night in *Bx* mutants. At ZT2, the *Bx* mutants showed reduced maximal axonal crosses relative to those in control flies that were not significantly different from those observed at ZT14 (Figure 5A, 5B, and 5C). Interestingly, when *Bx* was overexpressed in *miR-263b* expressing cells results were similar to those in the *miR-263b^**KO**^* flies (Figure 5A, 5B, and 5C). The rhythmic change in structural plasticity was abolished in the *Bx* mutants or Bx overexpression in DD (Figure 5D and 5E). These results support our hypothesis that *miR-263b* regulates the fasciculation-defasciculation state of the PDF termini via effects on *Bx* expression. In addition, activation of PDF cells with TrpA1 in the *Bx* mutant background resulted in defasciculated sLNv dorsal termini at ZT14 (Figure 5F and 5G).

**Figure 5.**
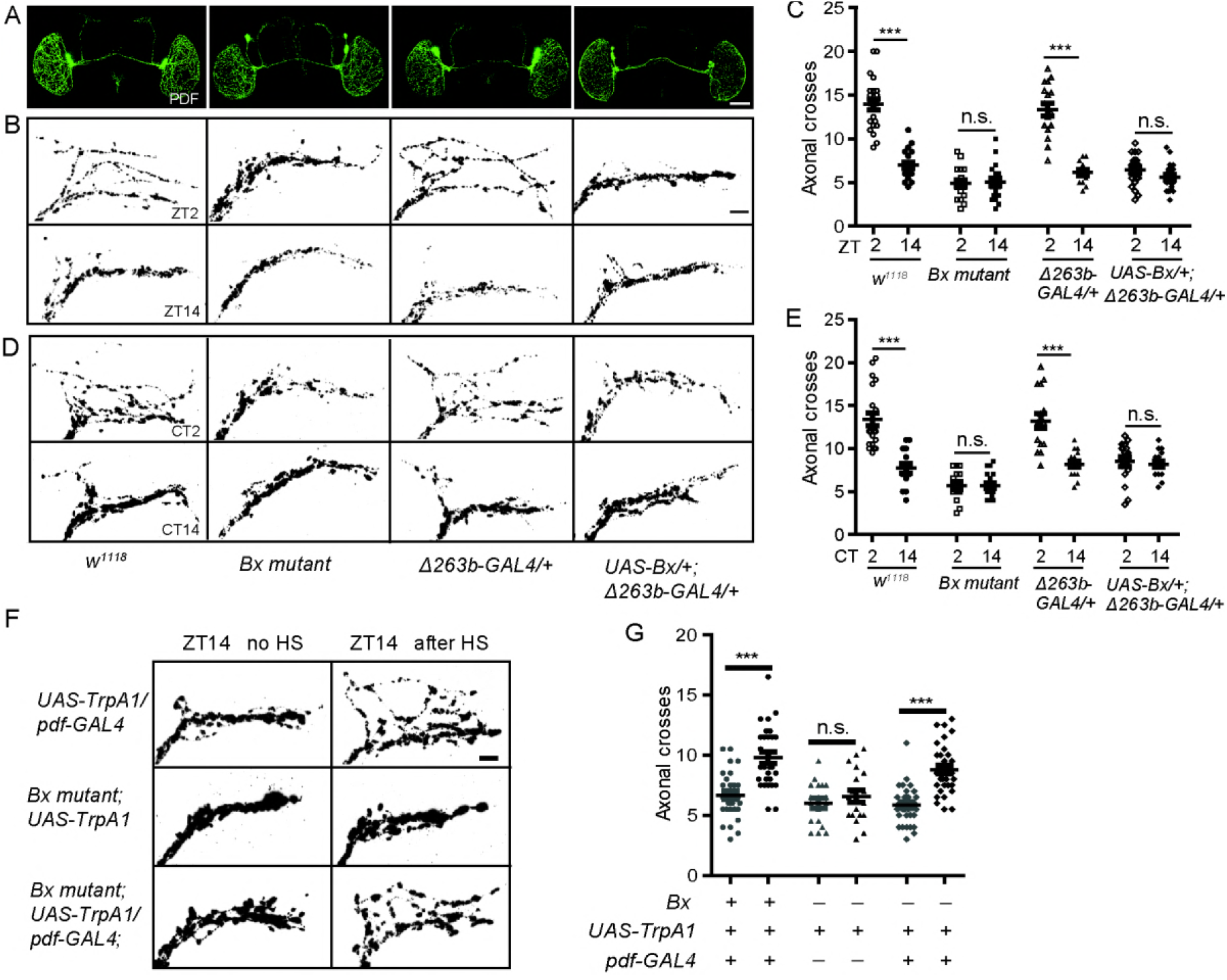
*Bx* regulates the arborization rhythms of sLNv dorsal projections. (A) Representative confocal images of brains of the indicated genotypes stained for PDF at ZT2. Scale bar, 75 μm. (B) Dorsal projections of sLNvs at ZT2 and ZT14. Flies were entrained for at least 3 days under LD conditions prior to the assay. Scale bar, 10 μm. (C) Quantification analysis of axonal morphology (fasciculation) of sLNv dorsal termini in LD conditions by Sholl’s analysis. Data represent means ± SEM (n=15–23). n.s. no significant. ***P< 0.001 determined by Student’s t test. (D) Representative confocal images of the projections at CT2 or CT14 during the second day of DD. Scale bar, 10 μm. (E) Quantification of axonal morphology of sLNv dorsal termini in DD conditions. Data represent means ± SEM (n=15–21). n.s. no significant. B***P< 0.001 determined by Student’s t test. (F) Confocal images of sLNv projections from flies stained with anti-PDF at ZT14. Conformation changes in response to 2-hour temperature elevation (HS) is shown on the right. The left show a control not subjected to temperature elevation. (G) Quantification of TrpA1-induced changes in axonal fasciculation. Data represent means ± SEM (n=19–34). n.s. no significant, ***P< 0.001 determined by Student’s t test.

### *miR-263b* binding sites in the *Bx* 3’UTR are essential for circadian function of Bx

To further confirm that *Bx* is a direct *miR-263b* target, we used the CRISPR/Cas9 system to mutate the potential *miR-263b* binding sites within the *Bx* 3’UTR in flies. The seed region of miRNAs (positions 2–7) are critical for activity (Brennecke, Stark, Russell, & Cohen, 2005). It has been shown that substitution of one nucleotide in the seed sequence is sufficient to reduce miRNA binding to the target mRNA (Brennecke et al., 2005). We recovered two fly lines (*Bx*\*1 and *Bx*\*2) with mutations in the *miR-263b* binding sites (Figure 6A). While *Bx*\*1 had a T to G point mutation in the seed sequences of one of the two *miR-263b* binding sites in the *Bx* 3’UTR, *Bx*\*2 had mutations in both sites. There were also several base changes near the seed sequence that differed in *Bx*\*1 and *Bx*\*2 (Figure 6A). We predicted that if *miR-263b* suppresses *Bx* abundance through the binding sites in the 3’UTR, *Bx*\*1 and *Bx*\*2 mutants should phenocopy *miR-263b^KO^*. As expected, both *Bx*\*1 and *Bx*\*2 mutants showed significantly impaired locomotor activity rhythms and reduced power of rhythms (Figure 6B-6E). The morning anticipation was also dramatically reduced or abolished, especially in the *Bx*\*2 mutants (Figure 6B and 6F). Furthermore, similar to the *miR-263b^KO^* flies, *Bx*\*2 mutants also had fasciculated sLNv projections and significantly decreased numbers of axonal crosses at ZT2 (Figure 6G and 6H). As flies with disrupted *miR-263b* binding sites within *Bx* 3’UTR show anatomical and behavioral phenotypes similar to those seen in *miR-263b^KO^* flies, we conclude that *miR-263b* regulates circadian rhythm by directly regulating *Bx* expression.

**Figure 6.**
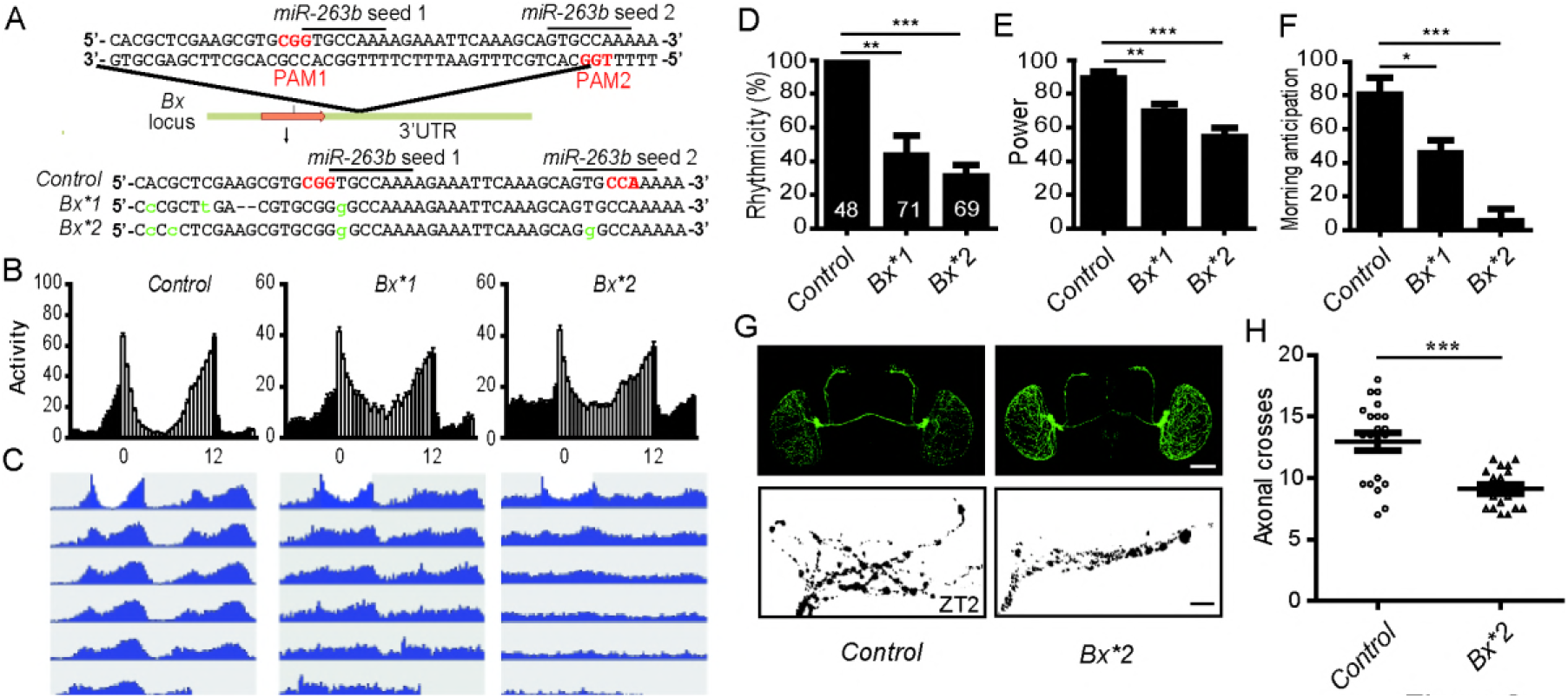
Mutation of the *miR-263b* binding site in the *Bx* 3’UTR leads to behavioral arrhythmicity. (A) Sequence in the *miR-263b* binding site region in the *Bx* 3’UTR and mutations induced using the CRISPR/Cas9 system. (B) Locomotor activity of adult male flies of indicated strains measured during 4 days of LD cycles and 5 days of DD. The white and black bars indicate day and night, respectively. (C) Actograms showing the average activities on the last day in LD and the 5 days in DD. Light represents the day and gray darkness. (D-F) Comparison of D) rhythmicity, E) power, and F) morning anticipation in indicated genotypes. The numbers of tested flies are shown in each column. Data represent means ± SEM (n=48–69). (G) Representative confocal images of control and *Bx*2* fly brains stained for PDF at ZT2. Scale bar, 75 μm for the whole-mount figure and 10 μm for the magnified images. (H) Quantification of axonal morphology from control and *Bx*2* flies. Data represent means ± SEM (n=18–21). *P<0.05, ^**^P<0.01, ***P< 0.001 determined by Student’s t test.

## Discussion

Here we demonstrate that *miR-263b* is critical for circadian behavior rhythms and axonal structural plasticity of pacemaker neurons. *miR-263b* modulates circadian rhythms via repression of *Bx*. Alterations in levels of *miR-263b* or *Bx* led to behavior arrhythmia and defects of arborization rhythm in sLNvs.

The mechanisms underlying the structural plasticity of sLNv dorsal projections are poorly understood. This plasticity is controlled by both circadian regulation and activity-dependent mechanisms. The maximal numbers of axon terminal branches are found in the early day and the minimal numbers in the night. This arborization rhythm exists even in constant darkness indicating this process is under circadian control. Manipulating the activity of sLNvs can change the axon plasticity, however. For example, electrical silencing the sLNvs during adulthood can reduce axon complexity (Depetris-Chauvin et al., 2011). Whether the circadian and activity regulatory mechanisms are independent of each other or interconnected is unknown. Recently, the transcription factor Mef-2 has been shown to control both circadian-and activity-dependent axonal fasciculation of sLNvs through the neural cell adhesion molecule Fas2 (Sivachenko et al., 2013). This indicates that the two mechanisms might be interconnected through Mef-2. Here we showed that *miR-263b* specifically regulates the circadian structural plasticity by suppressing *Bx* expression. Interestingly, the activity-dependent axonal changes are intact in *miR-263b* and *Bx* mutants. Thus, it is possible that different mechanisms influence circadian regulation and activity-dependent regulation of structural plasticity. It will be interesting to test whether there is a mechanism that only affects the activity-dependent structural changes in sLNvs.

Consistent with a report by Yang et al. (Yang et al., 2008), our data indicate that there is a cyclic expression of *miR-263b* in fly heads and that this expression is under clock control. Whereas the effect of *miR-263b* on structural plasticity occurred in the early morning, the maximal expression of *miR-263b* was late in the day and in early evening (Figure 1A). This apparent discrepancy between the expression peak and function peak may be due to the mechanism of miRNA action. Most miRNAs function through translational inhibition or degradation of target mRNAs so it is possible that the peak in protein abundance of miRNA targets is opposite the expression peak of the miRNA. In the future, it will be interesting to see whether *Bx* has a peak of abundance at early morning that matches its function.

Evidence suggests that *miR-263b* regulates *Bx* expression in the sLNv cell bodies. A previous study and data shown here clearly show that *Bx* is enriched in sLNvs (Fig S5, (Tsai et al., 2004)). That *Bx* is expressed in sLNvs suggests an autonomous mechanism since the *miR-263b* binding sites are critical. *miR-263b* is enriched in central nervous neurons (Yang et al., 2008); however, we cannot exclude a glial contribution of *miR-263b*. A recent study showed that glial expression of *miR-263b* is also required for circadian locomotor behavior (You, Fulga, Van Vactor, & Jackson, 2018). Tissue-specific knock out of *miR-263b* will help answering this question.

Here we demonstrated that *miR-263b* regulates *Drosophila* circadian locomotor rhythms and axonal structural plasticity of pacemaker neurons. A similar mechanism may exist in mammals. The suprachiasmatic nucleus (SCN) is the mammalian master circadian pacemaker (Mohawk, Green, & Takahashi, 2012; Reppert & Weaver, 2002). In SCN, a group of neurons expressing vasoactive intestinal peptide (VIP) receive retinal light input to synchronize the clock (Morin & Allen, 2006). VIP is the functional homologue of *Drosophila* PDF. Interestingly, circadian structural plasticity was found in the glutamatergic synapse on the VIP neurons, which maybe important for light entrainment (Girardet et al., 2010). As a highly conserved miRNA, *miR-263b* has a homologue in vertebrates: *miR-183*. Expression of *miR-183* is enriched in both retina and the pineal gland (Clokie, Lau, Kim, Coon, & Klein, 2012; Xu, Witmer, Lumayag, Kovacs, & Valle, 2007), Furthermore, in the rat pineal gland, *miR-183* is rhythmically expressed with a peak at around ZT12 (Clokie et al., 2012). This oscillation of *miR-183* suggests a potential circadian function. Bx is a *Drosophila* LIM-only protein. *In silico* miRNA target prediction with targetscan identified a highly conserved binding site of *miR-183* in the 3’UTR of *Lmo3*, a LIM-only protein conserved from rodents to primates (Dambal, Shah, Mihelich, & Nonn, 2015). Our results establish *miR-263b* as an important regulator of circadian locomotor behavior and suggest that a highly conserved miRNA-LIM-only protein pathway regulates circadian rhythms in organisms as diverse as flies and mammals.

## Materials and Methods

### Fly stocks

The following strains were used in this study: *w^1118^, Clk^Jrk^, y w; Pdf-GAL4/CyO, UAS-TrpA1, Δ263b-GAL4/TM6B* (Hilgers et al., 2010); *UAS-Bx* and *Bx* mutant (Bejarano, Smibert, & Lai, 2010); *Bx-GAL4, UAS-miR-263b, miR-263b^KO^* (obtained from the Bloomington Stock Center). All the flies were raised on standard cornmeal/agar medium at 25 °C under 12 hour:12 hour LD cycle.

### Behavioral experiments and analysis

Adult male flies (2–5 days old) were used to test locomotor activity rhythms. Flies were entrained for 4 days LD cycle at 25 °C, using about 500 lux light intensities, and then released into constant darkness (DD) at 25 °C for at least 5 days. Locomotor activity was recorded with a TriKinetics Activity Monitor in an I36-LL Percival Incubator. FAAS-X software was used to analyze behavioral data (Grima et al., 2002). Actograms were generated with a signal-processing toolbox for MATLAB. The morning anticipation amplitude was determined by assaying for the locomotor activity as described (Zhang & Emery, 2013).

### Whole-mount immunohistochemistry and quantification

Whole-mount immunohistochemistry of fly brains were done as previously described (Zhang et al., 2010). For PDF staining, adult flies were entrained to LD for 4 days and dissected at ZT2 or 14. For PER staining, flies were entrained to LD for 4 days and then release into DD. Brains were dissected on the second day of DD at six time points. Mouse anti-PDF (1:400), rabbit anti-GFP (1:600), rabbit anti-GFP (1:1500) antibodies were used. All samples were imaged using a Leica TCS SP8 confocal microscope with a constant laser setting for each time point. Eight to ten brains for each genotype were dissected for imaging. ImageJ software (Schneider, Rasband, & Eliceiri, 2012) was used for GFP, PER and PDF quantification from at least five brains. For quantification, the average signals of three neighboring background areas were subtracted from signal intensity in each circadian neuron. Each experiment was conducted three times.

### Quantitative real-time PCR

30–40 flies were collected at the indicated time points, heads were isolated on dry ice and stored at –80 °C. Total RNA, including miRNA, was purified with miRcute miRNA isolation kits (TIANGEN). Reverse transcription and real-time PCR of *miR-263b* and *2s rRNA* were performed with first-stand cDNA synthesis kits and miRcute miRNA qPCR detection kits (SYBR Green) (TIANGEN). For *miR-263b* the following primer was used: 5′GCGTTCTCCTTGGCACTGGG. *2s* was used for normalization and the following primer was used: 5′-TGCTTGGACTACATATGGTTGAGGG. Each experiment was conducted three times.

### S2 cell luciferase assay

The full-length 3’UTR of *Bx* and about 400 base pairs of coding region of *miR-263b* were amplified by PCR using primeSTAR HS DNA Polymerase (TaKaRa). The *Bx* 3’UTR was cloned into a pAc5.1-firefly luciferase-V5-His vector and the *miR-263b* coding region was cloned into a pAc5.1-V5-His vector (Invitrogen). The *miR-263b* seed-targeted sequence in the 3’UTR of *Bx* was mutated from GTGCCAA to CTACTCG using site-directed, ligase-independent mutagenesis (J. Chiu, March, Lee, & Tillett, 2004). We co-transfected 100 ng pAc-firefly luc-Bx 3’UTR (pAc-firefly luc-Bx 3’UTR mutant), 1 μg pAc-miR-263b, and 100 ng pAc-Renilla luc (transfection control) into S2 cells. Luciferase activity (Dual Luciferase System, Promega) was measured two days after transfection. Each experiment was conducted three times.

### Analysis of axonal morphology by modified Sholl’s method

The following fly genotypes were used: *pdf-GAL4/UAS-TrpA1* (control), *Bx mutant*; *UAS-TrpA1, Bx mutant; pdf-GAL4/UAS-TrpA1, UAS-TrpA1/*+; *miR-263b^KO^, pdf-GAL4/UAS-TrpA1*; *miR-263b^KO^*. For the analysis of activity-dependent changes in axonal morphology, *pdf-GAL4/UAS-TrpA1, Bx mutant; pdf-GAL4/UAS-TrpA1, pdf-GAL4/UAS-TrpA1; miR-263bKO, UAS-TrpA1; miR-263b^KO^, Bx mutant; UAS-TrpA1* flies were raised at 18 °C and entrained in LD cycles at 18 °C for at least 3 days, then shifted to 30 °C at ZT12. Flies were dissected 2 hours later (ZT14) and stained with anti-PDF. Structural plasticity was analyzed as reported (Sivachenko et al., 2013). Each experiment was conducted three times.

### Statistics analysis

Statistical analysis was performed with SPSS statistics 17.0. P values was obtained with t-test and considered n.s. no significant, significant at *P < 0.05 and extremely significant at *** P < 0.001.

## Acknowledgements

We thank Dr. Yi Liu and Dr. Patrick Emery for carefully reading the manuscript and suggesting improvements. We thank Dr. Steve Cohen for the *miR-263b^KO^* and *Δ263b-GAL4* fly strains and Dr. Eric Lai for the *Bx* mutant and *UAS-Bx* fly strains. We also thank Dr. Ralf Stanewsky for the anti-PER antibodies. We thank the Bloomington stock Center for various fly stocks. We also thank the Developmental Studies Hybridoma Bank for PDF antibodies. This work is supported by grants from the National Nature Science Foundation of China (Grant numbers 31572317 and 31730076) to Zhangwu Zhao and the China Scholarship Fund. Wenfeng Chen’s work is supported by the National Natural Science Foundation of China (grant number 31601894), and the Fujian Natural Science Foundation (grant. number 2017J0106). Yong Zhang’s lab is supported by the National Institutes of Health COBRE Grant P20 GM103650.

## Author Contributions

Z.Z. and Y.Z. supervised the project and designed the experiments. X.N., W.C., and W.B. performed the experiments and analysis. Y.Z., Z.Z., and X.N. wrote the manuscript.

---

**Figure S1.**
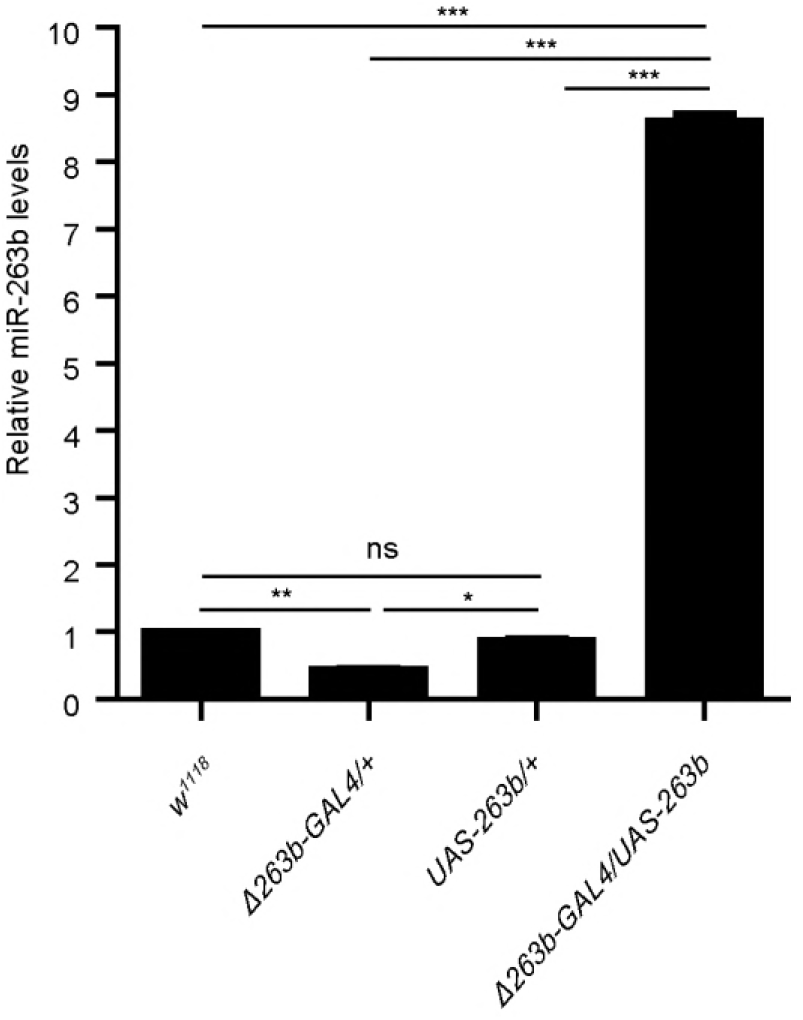
*miR-263b* expression level in relevant flies. Quantitative realtime PCR analysis of total RNA prepared from adult brains at ZT13. The relative expression levels were normalized to 2S RNA levels and were further normalized to *w*^1118^ control. Data represent mean ± SEM. n.s. no significant, *P<0.05, ^**^P<0.01, ***P< 0.001 determined by Student’s t test.

**Figure S2.**
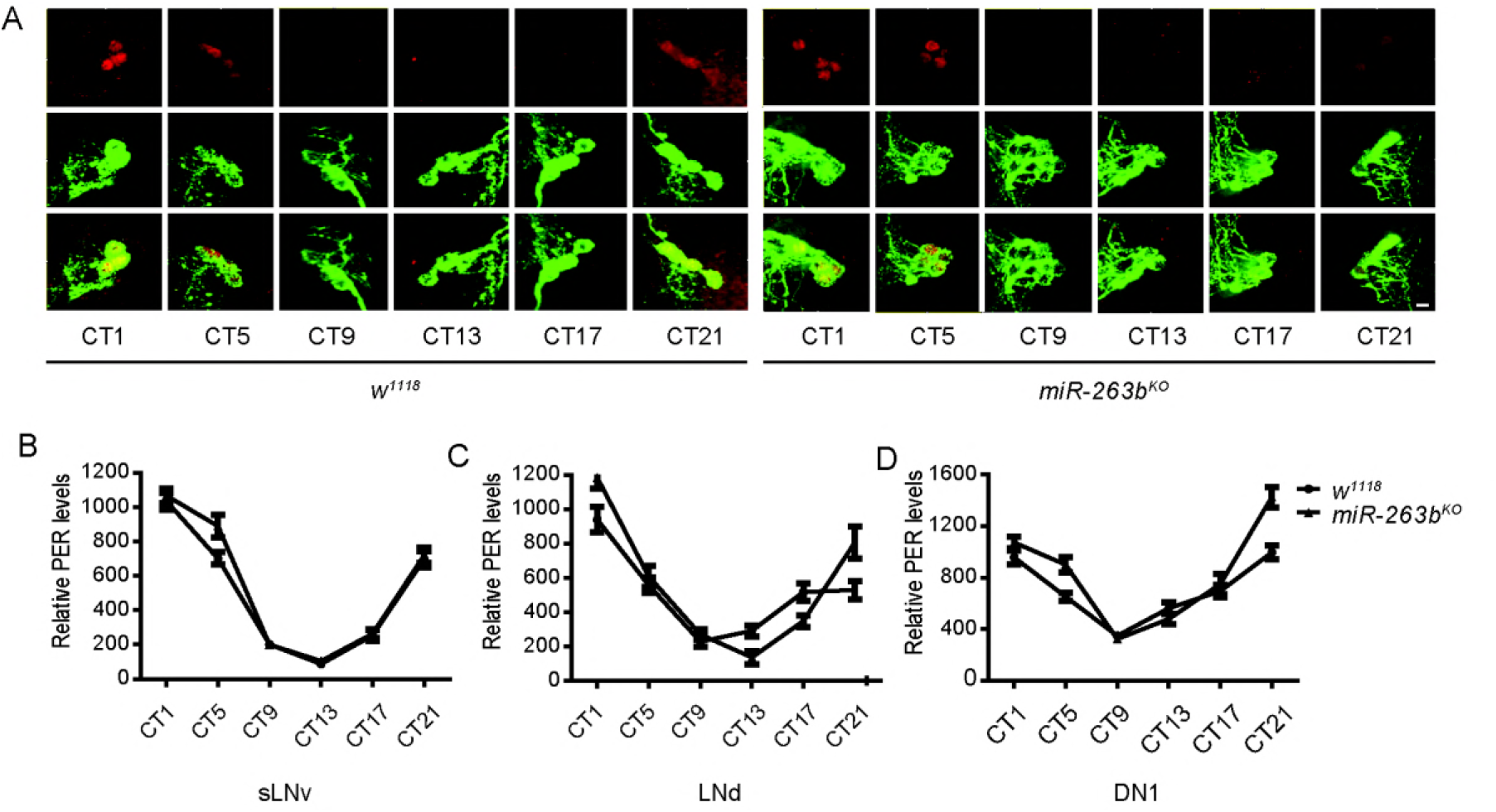
The molecular pacemaker is not affected in miR-263b^KO^ flies. (A) Representative confocal images of sLNvs from *w*^1118^ and *miR-263b^KO^* flies dissected at six time points (circadian time, CT) during the second day of DD and stained with anti-PDF (green) and anti-PER (red) antibodies. Scale bar, 10 µm. (B-D) Quantification of PER staining in sLNvs, LNds, and DN1s at different circadian time points. Data represent mean ± SEM (16–19).

**Figure S3.**
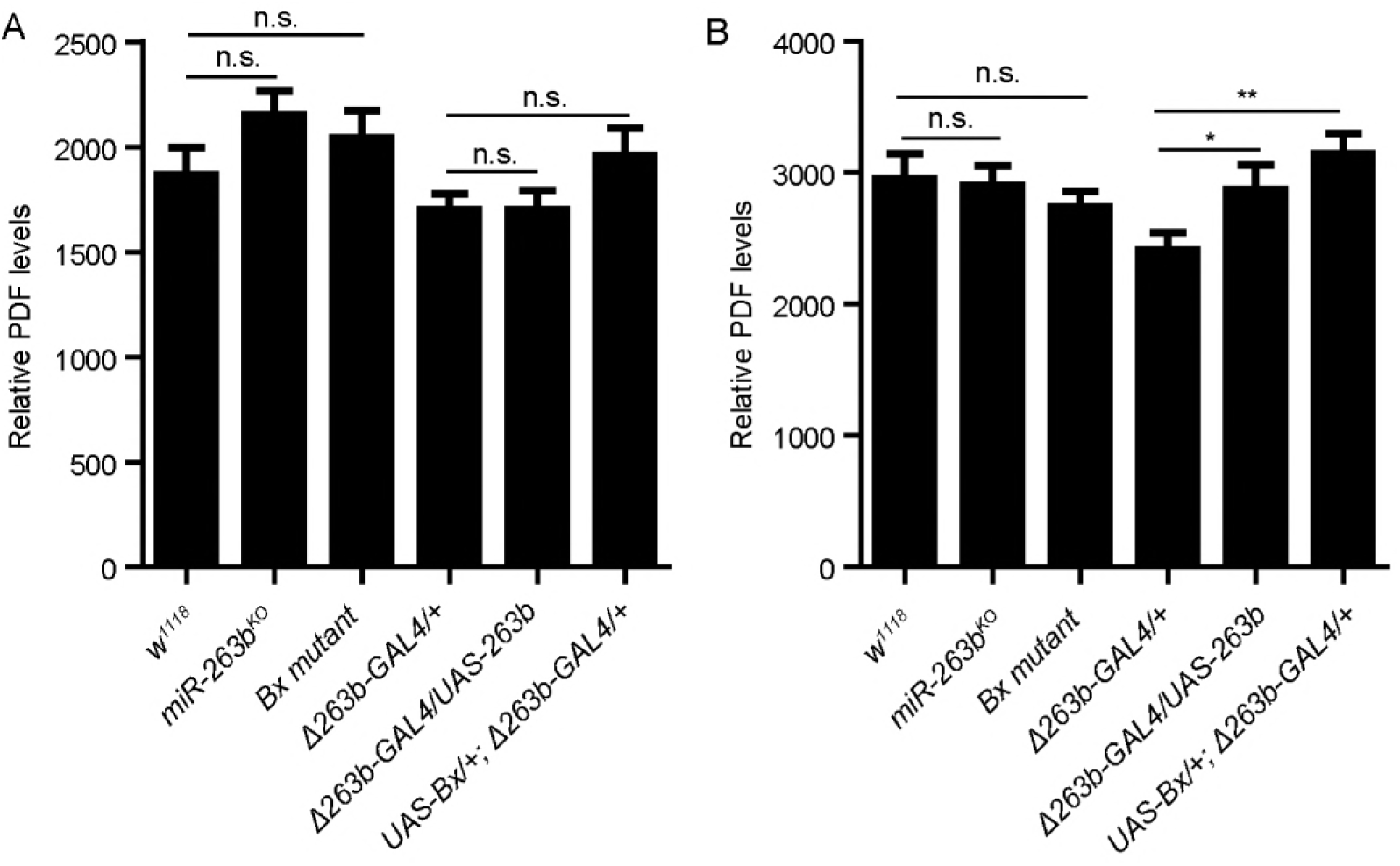
PDF levels in sLNvs soma are not significantly changed under LD conditions. (A-B) Quantification of PDF staining in sLNvs at ZT2 and ZT14. Data represent mean ± SEM (n=18–21). n.s. no significant, *P<0.05, ^**^P<0.01, ***P< 0.001 determined by Student’s t test.

**Figure S4.**
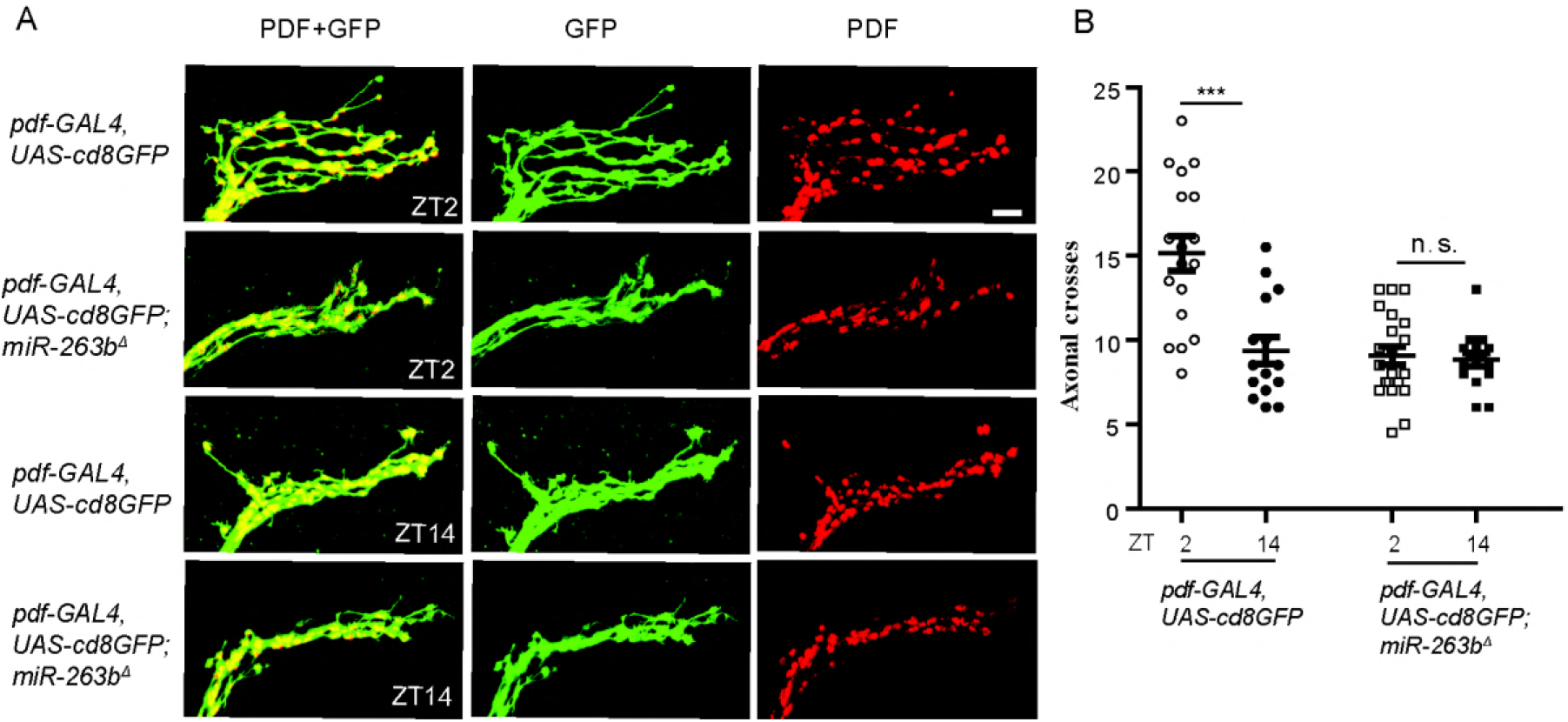
shows the axonal projection of an sLNv labeled by a membrane-tethered GFP. (A) Representative confocal images of fly brains stained for GFP (green) and PDF (red) dissected at ZT2 and ZT14. Scale bar, 10 µm. (B) Quantification analysis of axonal morphology of sLNv dorsal termini. Data represent mean ± SEM (n=15–21). n.s. no significant, ***P< 0.001 determined by Student’s t test.

**Figure S5.**
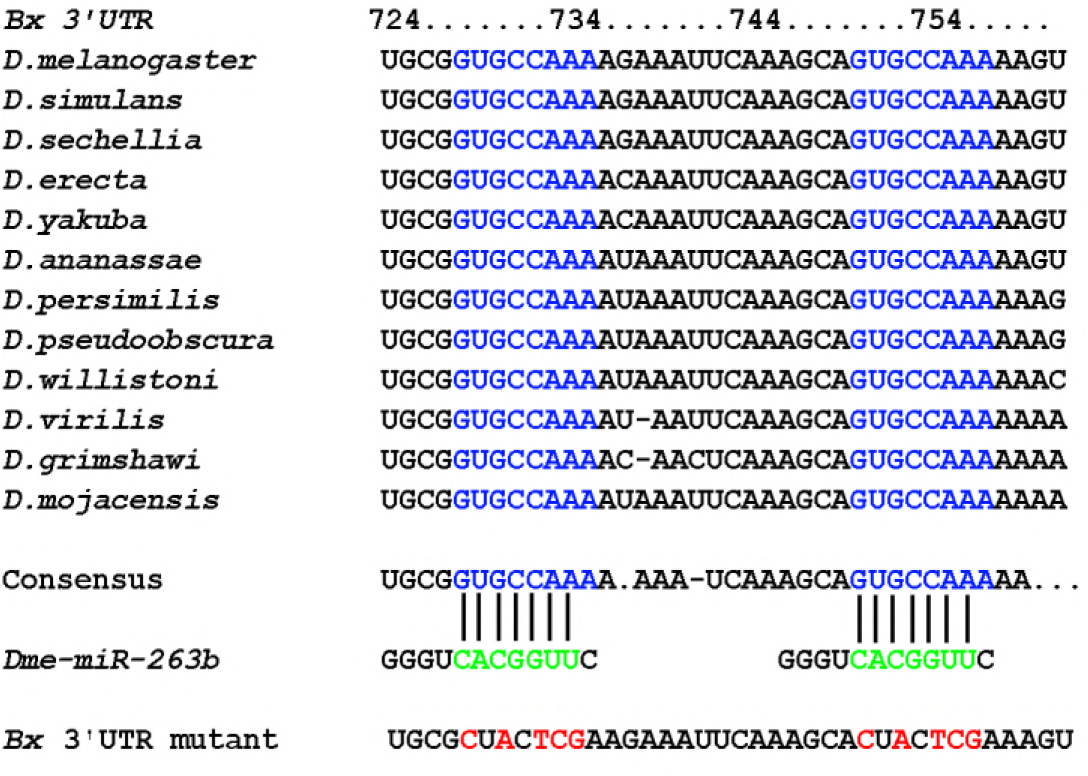
Predicted *miR-263b* binding site conservation in the *Bx* 3’UTR among *Drosophila* species. Blue letters indicate conserved sequences. Green letters indicate miR-263b seed region; red letters indicate positions of mutations in *Bx*3’UTR made for S2 cell reporter gene assay.

**Figure S6:**
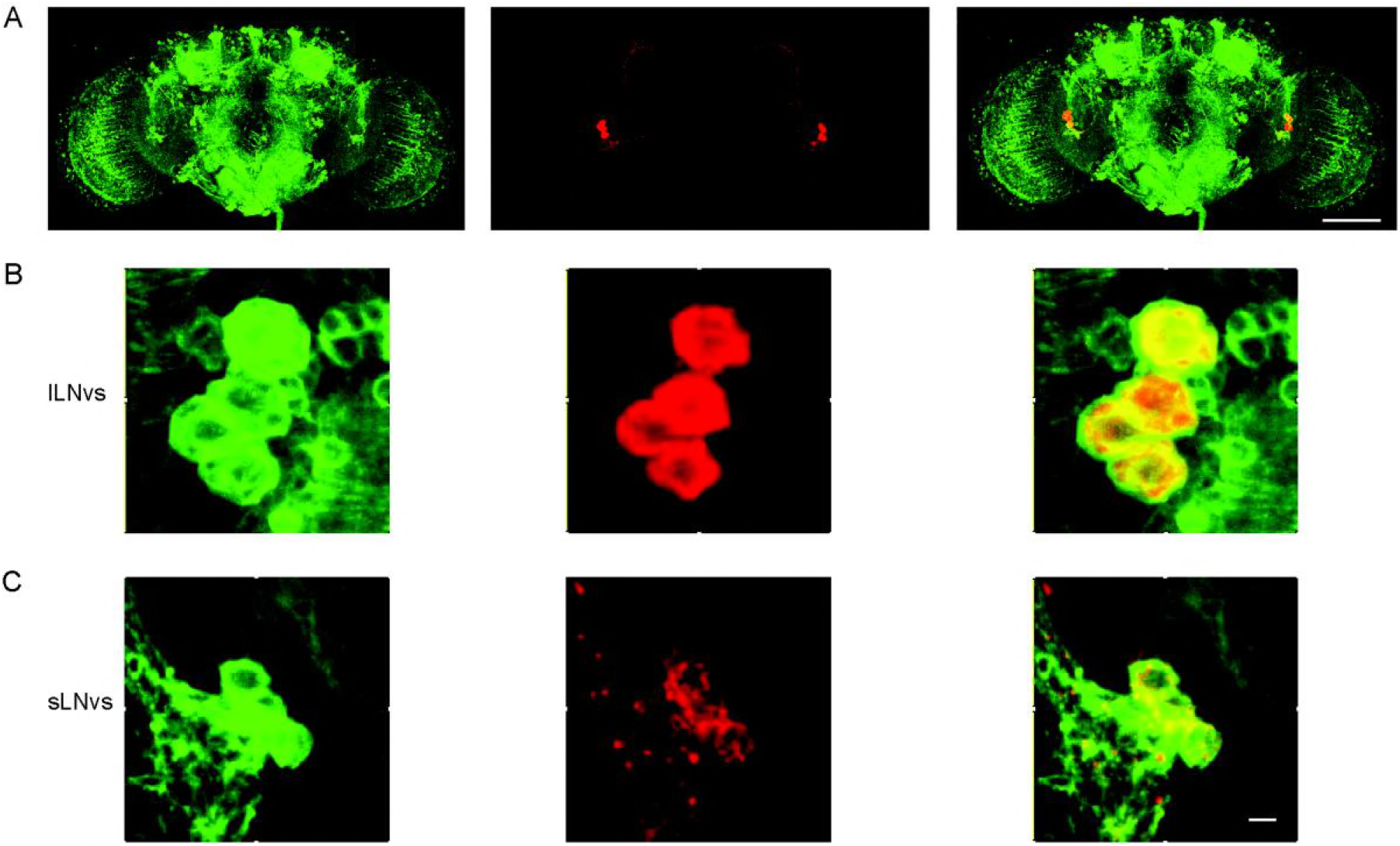
*Bx* is expressed in the PDF positive sLNvs and lLNvs. (A-C) Representative confocal images of fly brains, lLNvs and sLNvs stained for GFP (green) and PDF (red) dissected at ZT13. Scale bar, 100 μm.

